# OracleScreen-LILRB4: Machine Learning-Guided Discovery of Myeloid Immune Checkpoint Binders Validated in Patient-Derived Cells

**DOI:** 10.64898/2026.06.17.732859

**Authors:** Somaya A. Abdel-Rahman, Moustafa T. Gabr

## Abstract

The identification of small molecule modulators of immune checkpoint proteins remains a significant challenge in drug discovery due to the flat, featureless nature of protein-protein interaction interfaces and the characteristically low hit rates observed in conventional high-throughput screening campaigns. Here we report OracleScreen-LILRB4, an ensemble machine learning framework trained on quantitative biophysical screening data from two structurally diverse compound libraries (19,800 compounds total) screened against the myeloid immune checkpoint leukocyte immunoglobulin-like receptor B4 (LILRB4/ILT3). By formulating binding prediction as a regression task targeting continuous ΔF_norm_ values rather than binary hit classifications, OracleScreen-LILRB4 achieved a mean Spearman R of 0.61 and ROC-AUC of 0.86 under scaffold-aware cross-validation. Prospective virtual screening of a 45,760-member compound library and experimental validation of the top 200 predictions yielded a 28.5% hit rate, representing a 15.0-fold enrichment over baseline, with 16 compounds demonstrating nanomolar-affinity LILRB4 (ILT3) engagement. Lead compounds **ORS-22** and **ORS-14** restored anti-tumor immune activity across patient-derived colorectal cancer and acute myeloid leukemia co-culture systems, reversing SCG2-mediated immunosuppression and recovering cytotoxic T-cell function. These findings establish OracleScreen-LILRB4 as an effective computational framework for accelerating small molecule discovery against non-enzymatic immune checkpoint targets.

**Graphical abstract:** 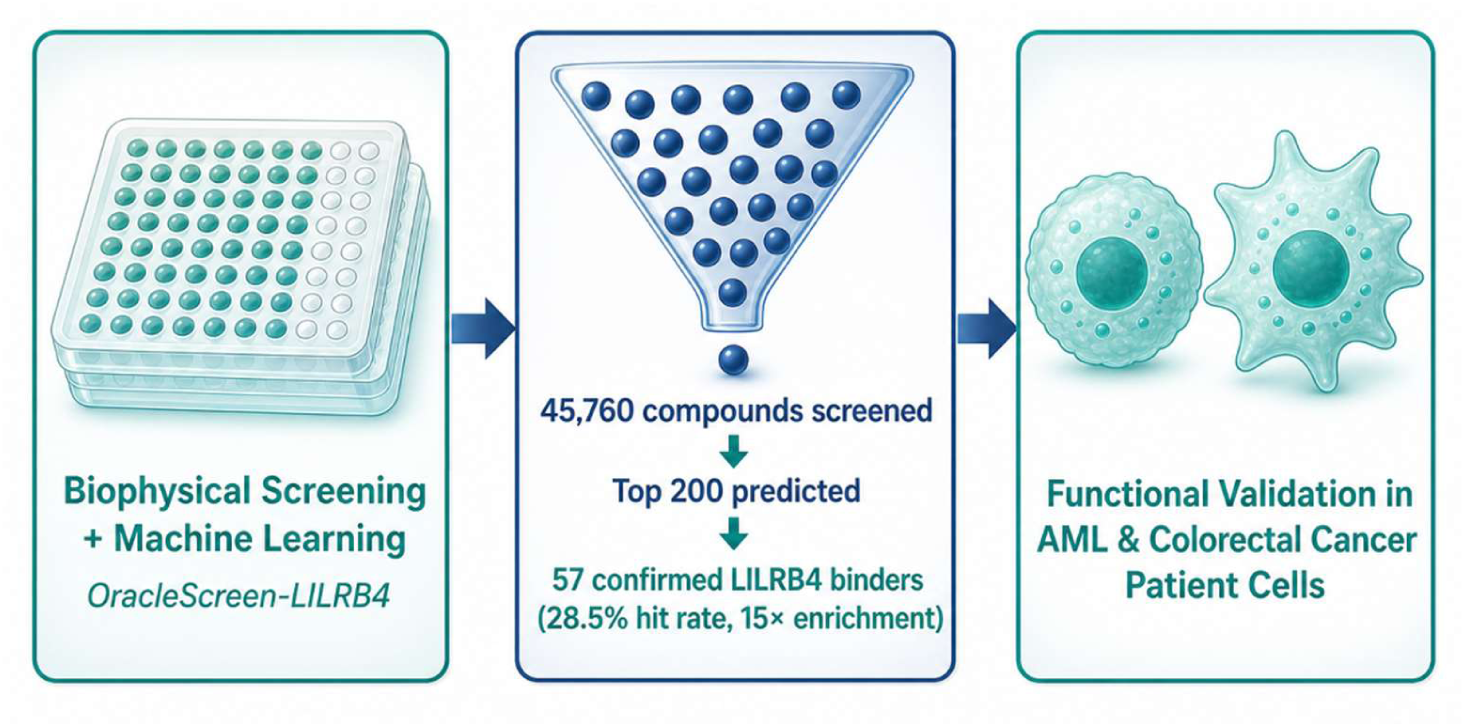

---

High-throughput screening (HTS) continues to serve as a foundational platform in small molecule drug discovery; however, its practical efficiency remains limited. Even under optimized conditions, screening campaigns commonly generate hit rates of only 1-2%, and for difficult biological targets, success rates often fall below 0.3%, while simultaneously demanding substantial investments in reagents, purified protein, instrumentation, and operational time.^1,2^ These limitations become particularly evident for immune checkpoint proteins, which include inhibitory receptors as well as stimulatory receptors.^3,4^ Immune checkpoints are critical regulators of T cell function and anticancer immunity, and their clinical relevance has been firmly established. The approval of relatlimab combined with nivolumab for dual LAG-3/PD-1 blockade in metastatic melanoma,^5^ together with ongoing therapeutic development programs directed toward TIGIT,^6^ ICOS,^7^ and CD28,^8^ highlights the significant translational potential of this target space. Nevertheless, most immune checkpoint proteins signal through protein-protein interaction (PPI) interfaces that are structurally broad, relatively flat, and poorly defined, making them exceptionally difficult to modulate using traditional small molecules.^9,10^ As a result, experimental screening campaigns against these targets frequently produce extremely poor hit rates.^11^

Machine learning approaches have increasingly been integrated into modern discovery pipelines to overcome these bottlenecks by enabling computational prioritization across ultra-large chemical libraries.^12–14^ Among the most widely adopted molecular representations, Morgan fingerprints capture local atomic environments through circular substructure encoding and have consistently demonstrated strong predictive performance when combined with physicochemical descriptors in ensemble modeling frameworks.^15,16^ Random forest, gradient boosting, and related tree-based ensemble methods are particularly well-suited to high-throughput screening datasets, where class imbalance, structural diversity, and moderate dataset sizes present challenges for deep learning architectures that require substantially larger training corpora. Scaffold-aware cross-validation strategies have further strengthened the reliability of such models by ensuring that chemically distinct scaffolds are partitioned across training and test folds, providing a more stringent and generalizable assessment of predictive performance than conventional random splitting approaches.^15,16^

Among newly recognized myeloid immune checkpoints, leukocyte immunoglobulin-like receptor B4 (LILRB4), also referred to as ILT3, has emerged as an important mediator of immunosuppression in both solid malignancies and hematologic cancers.^17–21^ LILRB4 is abundantly expressed in immunosuppressive myeloid compartments, including tumor-associated macrophages (TAMs), myeloid-derived suppressor cells (MDSCs), dendritic cells, and monocytic leukemia populations, where it contributes to the establishment of tolerogenic immune states and inhibition of T-cell function.^22–25^ Engagement of LILRB4 activates intracellular immunoreceptor tyrosine-based inhibitory motifs (ITIMs), triggering recruitment of the phosphatases SHP1 and SHP2 and ultimately dampening inflammatory and cytotoxic signaling pathways.^26–28^ Increased LILRB4 expression has been correlated with unfavorable clinical outcomes, impaired T-cell responses, resistance to immune checkpoint therapies, and accelerated tumor progression in multiple cancer settings.^29–31^ Antibody-based blockade of LILRB4 in preclinical models has been shown to restore T-cell activity, reprogram suppressive myeloid cell populations, and enhance antitumor immune responses, collectively supporting LILRB4 as a compelling therapeutic target in cancer immunotherapy.^32–34^ Nevertheless, most LILRB4-directed therapeutic strategies to date have focused on biologic modalities, whereas the development of small molecule LILRB4 modulators remains comparatively limited.

In this study, we report the development of OracleScreen-LILRB4, an ensemble machine learning framework trained on biophysical high-throughput screening data, specifically designed to address the small molecule discovery gap for the myeloid immune checkpoint LILRB4. Using Dianthus biophysical screening of two independent compound libraries, we identified LILRB4 binders and applied OracleScreen-LILRB4 to virtually screen an additional large compound library, prioritizing top candidates for experimental validation. Confirmed hits were subsequently evaluated in patient-derived acute myeloid leukemia (AML) cells, establishing their functional relevance in a clinically representative cellular context. Collectively, these findings demonstrate the utility of biophysical screening-trained machine learning for accelerating small molecule discovery against non-enzymatic myeloid immune checkpoints.

To enable machine learning-guided discovery of LILRB4 small molecule binders, we first established a biophysical training dataset by screening two structurally diverse compound libraries against recombinant human LILRB4 extracellular domain using Dianthus-based TRIC technology (Figure 1A). Screening of the Enamine Protein-Protein Interaction Library (15,000 compounds) identified 182 direct LILRB4 binders (1.21% hit rate, Figure 1B), while screening of the Enamine Immunology-Focused Library (4,800 compounds) yielded 104 additional binders (2.17% hit rate, Figure 1C), for a combined training dataset of 19,800 compounds comprising 286 confirmed LILRB4 binders. Hits were defined as compounds producing ΔF_norm_ responses exceeding three standard deviations above vehicle-treated controls.

**Figure 1.**
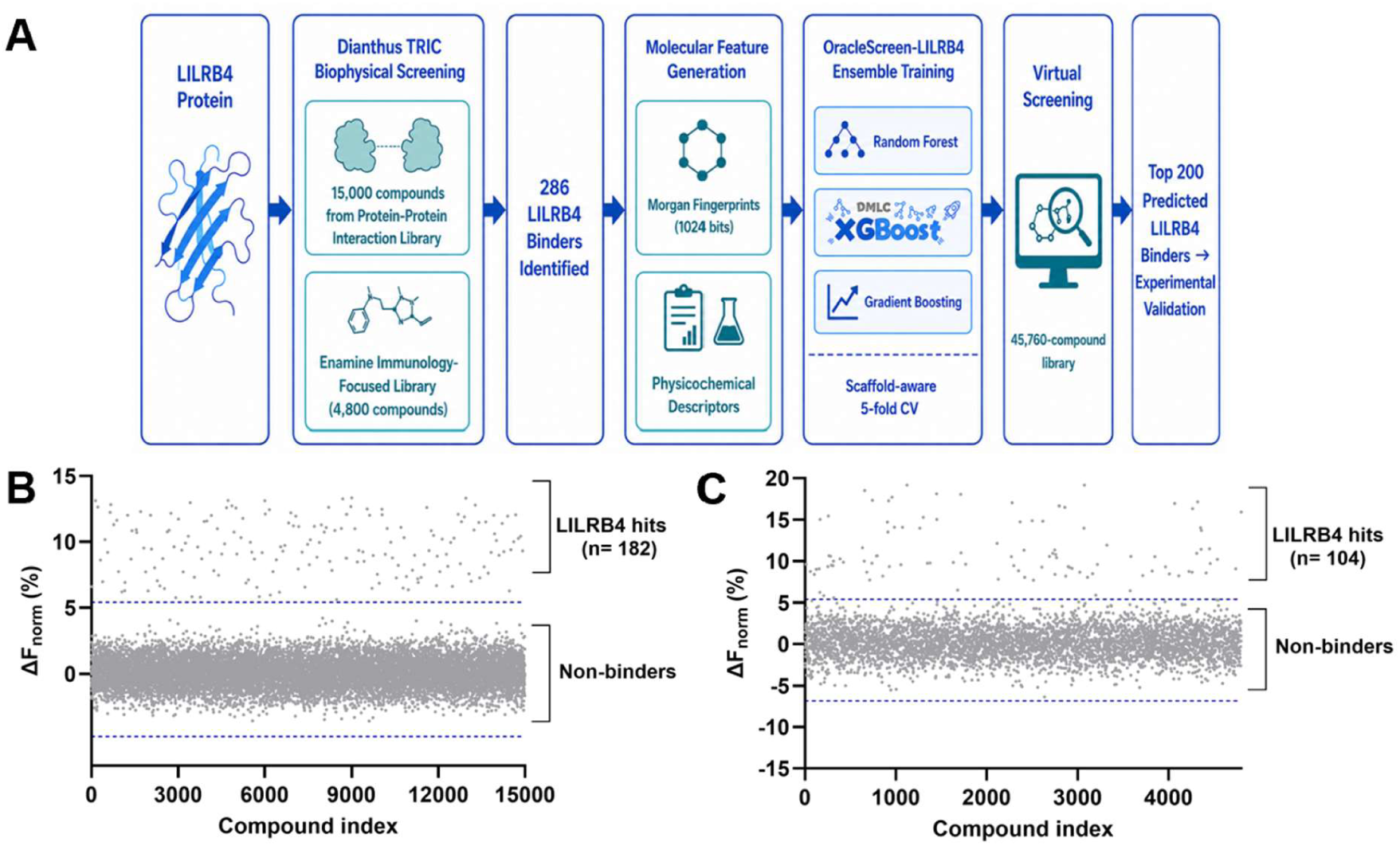
OracleScreen-LILRB4 machine learning pipeline and primary biophysical screening results. **(A)** Schematic overview of the OracleScreen-LILRB4 workflow. LILRB4 extracellular domain was screened against two compound libraries using Dianthus-based biophysical screening. Confirmed binders were used to train an ensemble machine learning framework integrating Morgan fingerprint representations with physicochemical descriptors under a scaffold-aware five-fold cross-validation strategy. The trained model was applied to virtually screen a 45,760-member compound library, and the top 200 predicted binders were selected for experimental validation. **(B)** Primary Dianthus/TRIC screening results for the Enamine Protein-Protein Interaction Library (15,00 compounds). Compounds exceeding the hit threshold of mean + 3SD are designated as LILRB4 binders (n = 182). **(C)** Primary Dianthus/TRIC screening results for the Enamine Immunology-Focused Library (4,800 compounds). Compounds exceeding the hit threshold are designated as LILRB4 binders (n = 104).

The combined dataset of 19,800 compounds with experimentally measured ΔF_norm_ binding signals was used to train OracleScreen-LILRB4, an ensemble machine learning framework designed to predict quantitative LILRB4 binding propensity directly from molecular structure. Rather than treating the screening output as a binary classification problem, OracleScreen-LILRB4 was formulated as a regression task targeting continuous ΔF_norm_ values, thereby preserving the full quantitative information content of the biophysical assay and avoiding information loss inherent to threshold-based hit calling. Molecular representations were constructed by concatenating extended-connectivity Morgan fingerprints (radius = 2, 1024 bits) with ten physicochemical descriptors encompassing molecular weight, lipophilicity, hydrogen bond donor and acceptor counts, topological polar surface area, rotatable bond count, aromatic ring count, fraction of sp3 carbons, heteroatom count, and ring count. This hybrid representation strategy captures both local atomic environment information and global drug-like properties relevant to protein binding. Three complementary ensemble algorithms were implemented, Random Forest, XGBoost, and Gradient Boosting, and their predictions were averaged to generate a consensus ensemble score, reducing model-specific variance and improving robustness of rank-ordering across chemically diverse compound libraries.

Model performance was rigorously evaluated using a scaffold-aware five-fold cross-validation strategy, in which compounds sharing identical Murcko scaffolds were assigned to the same fold. This approach enforces a more stringent generalization assessment than random splitting by ensuring that the model is evaluated exclusively on structurally novel scaffolds absent from the training set, directly simulating the prospective virtual screening scenario. The ensemble model achieved a mean Spearman R of 0.61 across five cross-validation folds (Random Forest: 0.57 ± 0.04; XGBoost: 0.63 ± 0.03; Gradient Boosting: 0.59 ± 0.05), demonstrating meaningful rank-order correlation between ensemble-predicted and experimentally measured ΔF_norm_ binding signals (Figure 2A). Consistent with the regression-based formulation, ensemble-predicted ΔF_norm_ scores also demonstrated strong discriminative performance between confirmed binders and non-binders, yielding a mean ROC-AUC of 0.86 (Random Forest: 0.82 ± 0.03; XGBoost: 0.89 ± 0.02; Gradient Boosting: 0.84 ± 0.03), despite the challenging class imbalance inherent to primary HTS datasets (1.44% overall hit rate) (Figure 2B). These findings confirm that quantitative binding signal prediction translates directly into accurate rank-ordering of active versus inactive compounds. The trained OracleScreen-LILRB4 ensemble was subsequently applied to virtually screen a 45,760-member compound library, assigning each compound a predicted ΔF_norm_ score reflecting its estimated LILRB4 binding signal. The resulting score distribution revealed a subset of high-scoring compounds well-separated from the bulk of the library at a predicted ΔF_norm_ cutoff of 8.4%, and the top 200 predicted binders were selected for experimental validation (Figure 2C). To interpret the structural determinants driving model predictions, SHAP analysis was performed on the Random Forest component. cLogP and aromatic ring count emerged as the most influential physicochemical features, suggesting that LILRB4 binding is preferentially associated with moderately lipophilic, aromatic-rich scaffolds capable of engaging hydrophobic surface patches and shallow protein-protein interaction interfaces, consistent with the hydrophobic and electrostatic character of the LILRB4 extracellular domain binding interface (Figure 2D).

**Figure 2.**
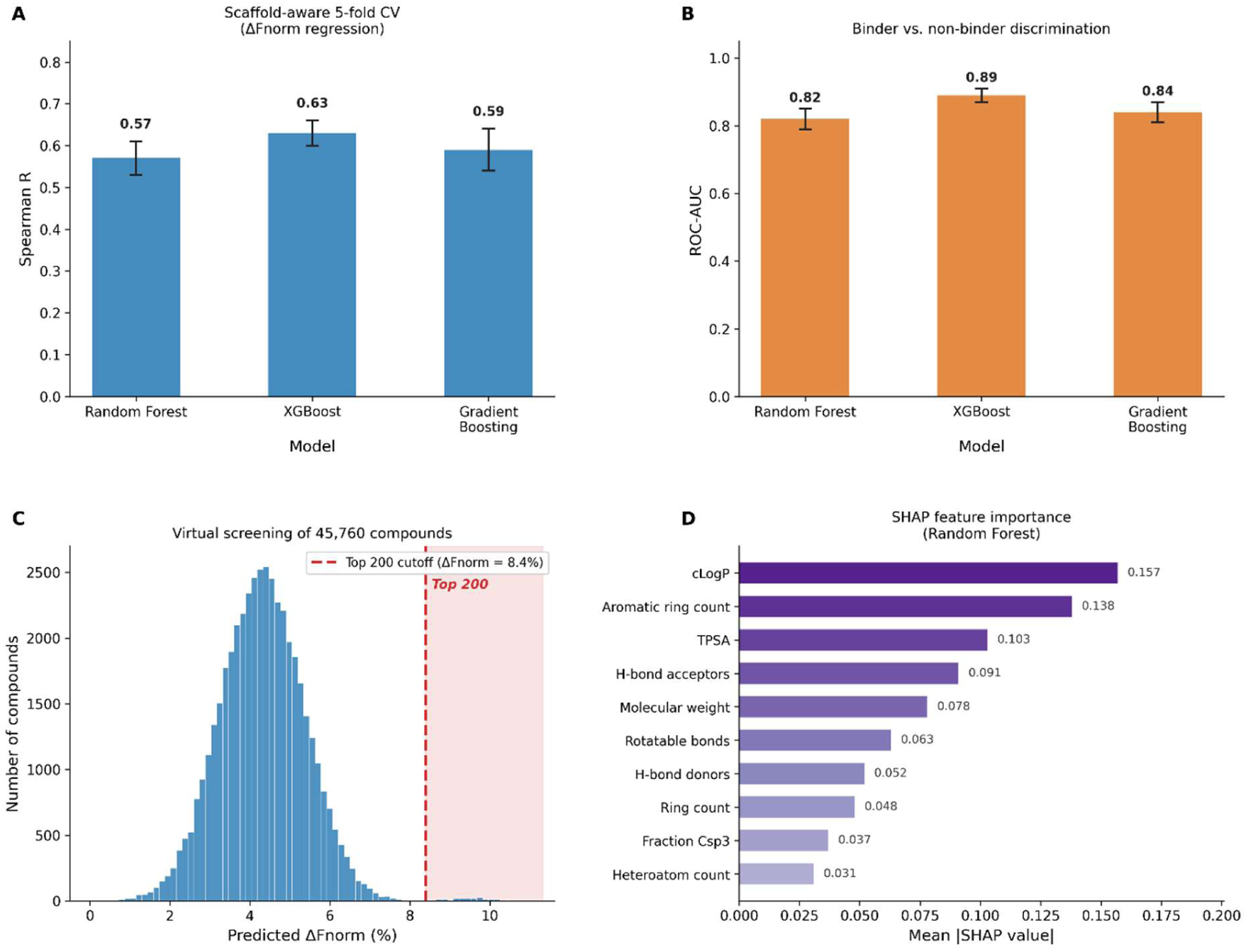
OracleScreen-LILRB4 model performance and virtual screening results. **(A)** Scaffold-aware five-fold cross-validation performance of the three ensemble model components evaluated by Spearman rank correlation between predicted and experimentally measured ΔF_norm_ binding signals. Data represent mean ± SD across five folds. **(B)** Discriminative performance of ensemble-predicted ΔF_norm_ scores for classification of confirmed LILRB4 binders versus non-binders, expressed as ROC-AUC. Data represent mean ± SD across five folds. **(C)** Virtual screening score distribution for the 45,760-member compound library. Each compound was assigned an ensemble-predicted ΔF_norm_ score reflecting estimated LILRB4 binding propensity. The dashed red line indicates the top 200 cutoff (predicted ΔF_norm_ = 8.4%); the shaded region denotes compounds selected for experimental validation. **(D)** SHAP feature importance analysis of the Random Forest model component. Mean absolute SHAP values are shown for all ten physicochemical descriptors, with cLogP and aromatic ring count identified as the most influential features driving LILRB4 binding predictions.

To prospectively validate the predictive performance of OracleScreen-LILRB4, the top 200 ML-prioritized compounds were purchased from Enamine and screened against recombinant human LILRB4 extracellular domain using Dianthus-based TRIC technology under identical conditions to the primary screening campaigns. Single-dose screening identified 57 confirmed LILRB4 binders, corresponding to a hit rate of 28.5%, representing a 15.0-fold enrichment over the combined baseline screening hit rate of 1.44% (Figure 3A). This prospective validation demonstrates that OracleScreen-LILRB4 successfully prioritized structurally novel LILRB4 binders from a large compound library, confirming the practical utility of biophysical screening-trained machine learning for target-focused compound enrichment.

**Figure 3.**
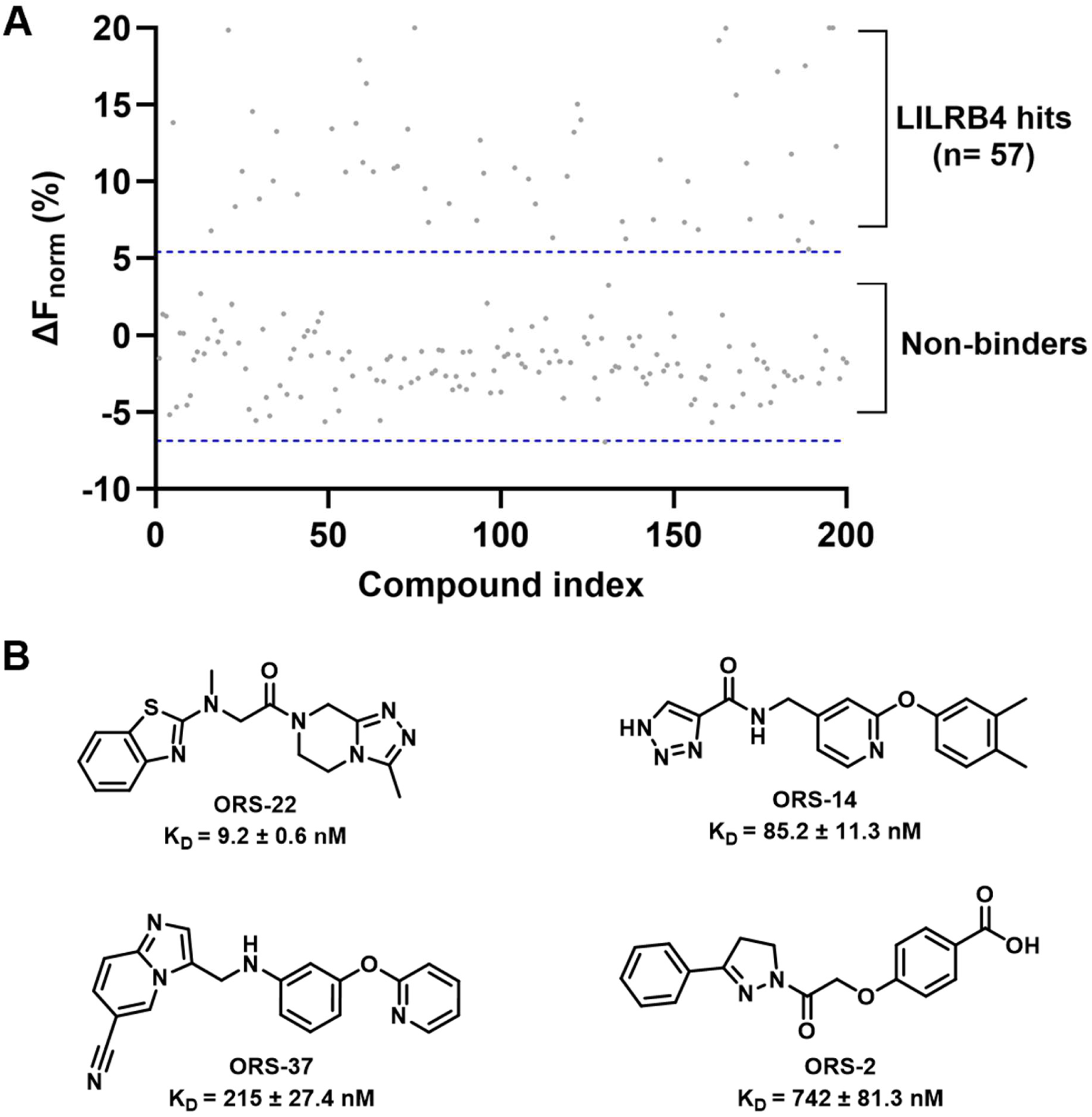
Prospective experimental validation of OracleScreen-LILRB4 predictions. **(A)** Primary Dianthus/TRIC screening results for the top 200 ML-prioritized compounds. Compounds exceeding the hit threshold of mean + 3SD are designated as LILRB4 binders (n = 57). **(B)** Chemical structures and equilibrium dissociation constants (K_D_) of the four highest-affinity LILRB4 binders identified from the ML-prioritized set. K_D_ values were determined by concentration-dependent Dianthus TRIC screening. All 16 confirmed binders with K_D_ values are reported in Table S1.

To quantitatively characterize binding affinities, the 57 confirmed hits were subjected to concentration-dependent Dianthus screening, revealing 16 compounds (Table S1) with clear dose-dependent binding profiles. Equilibrium dissociation constants (K_D_) were determined for all 16 compounds and are reported in Table S1. The four highest-affinity binders, designated **ORS-22**, **ORS-14**, **ORS-37**, and **ORS-2**, demonstrated K_D_ values of 9.2 ± 0.6 nM, 85.2 ± 11.3 nM, 215 ± 27.4 nM, and 742 ± 81.3 nM, respectively, establishing direct nanomolar-affinity engagement of the LILRB4 extracellular domain (Figure 3B). These compounds represent structurally diverse LILRB4 binders with drug-like physicochemical profiles, consistent with the cLogP and aromatic ring count features identified by SHAP analysis as key drivers of LILRB4 binding propensity.

Orthogonal biophysical and cellular target engagement studies further established **ORS-22** as the lead LILRB4-targeting compound. Concentration-dependent MST/TRIC analysis demonstrated direct binding of **ORS-22** to the recombinant LILRB4 extracellular domain with a K_D_ of 9.2 ± 0.6 nM, consistent with high-affinity and saturable ligand–protein interaction behavior (Figure 4A). The low nanomolar affinity observed for **ORS-22** positions it as the most potent small molecule LILRB4 binders identified in this study.

**Figure 4.**
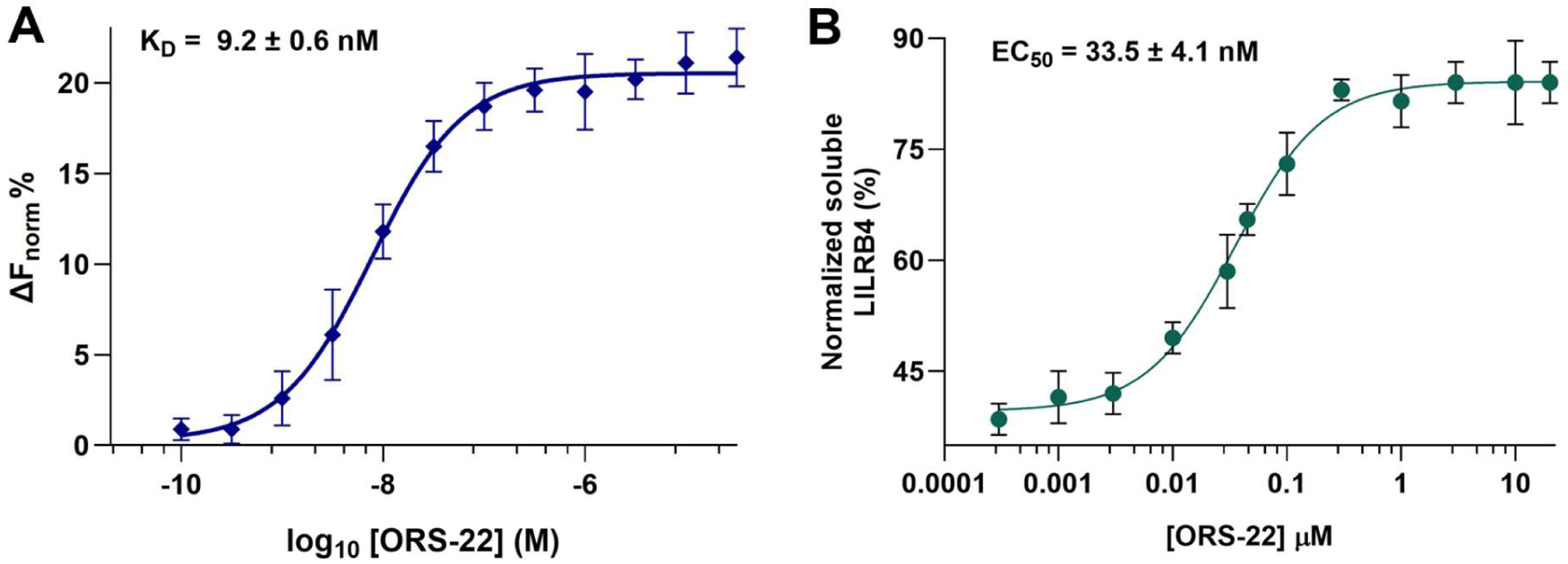
Orthogonal biophysical and cellular target engagement validation of ORS-22 against LILRB4. **(A)** Concentration-dependent MST/TRIC analysis demonstrating direct binding of **ORS-22** to the recombinant human LILRB4 extracellular domain. Data were fitted using a one-site binding model, yielding a K_D_ of 9.2 ± 0.6 nM. Data represent mean ± SD, n = 5. **(B**) CETSA demonstrating dose-dependent stabilization of LILRB4 following treatment with **ORS-22**, confirming cellular target engagement under physiologically relevant conditions. EC_50_ value was determined by nonlinear regression analysis. Data represent mean ± SD, n = 5.

To independently validate cellular target engagement using an orthogonal assay platform, a cellular thermal shift assay (CETSA) was performed. CETSA measures ligand-induced thermal stabilization of proteins within cells and is widely used to confirm direct compound–target engagement under physiologically relevant conditions. Treatment with **ORS-22** produced a robust dose-dependent increase in thermally stabilized soluble LILRB4, yielding an EC_50_ of 33.5 ± 4.1 nM (Figure 4B). The close agreement between the biochemical binding affinity and cellular stabilization potency strongly supports direct engagement of LILRB4 by **ORS-22** in both recombinant and cellular settings. Together, these orthogonal validation studies identify **ORS-22** as a potent and cell-active LILRB4 binder suitable for downstream functional characterization.

To determine whether direct pharmacological targeting of LILRB4 translates into functional restoration of anti-tumor immunity, the two highest-affinity compounds identified in this study, **ORS-22** and **ORS-14** (the two most potent compounds from this study), were evaluated in clinically relevant ex vivo co-culture systems representing both solid and hematologic malignancies (Figure 5A). In colorectal cancer co-cultures containing patient-derived PBMCs and HCT116 tumor cells, SCG2-mediated activation of LILRB4 induced a strongly suppressive immune phenotype characterized by reduced IFN-γ and IL-2 secretion together with increased tumor cell viability. Treatment with **ORS-22** or **ORS-14** (at 1 μM) significantly reversed these suppressive effects, restoring cytokine production and reducing tumor cell viability in a manner consistent with recovery of immune-mediated tumor killing (Figure 5B-D). Notably, **ORS-22** consistently demonstrated greater functional activity across multiple readouts, further supporting its prioritization as the lead compound emerging from the screening and biophysical validation workflow.

**Figure 5.**
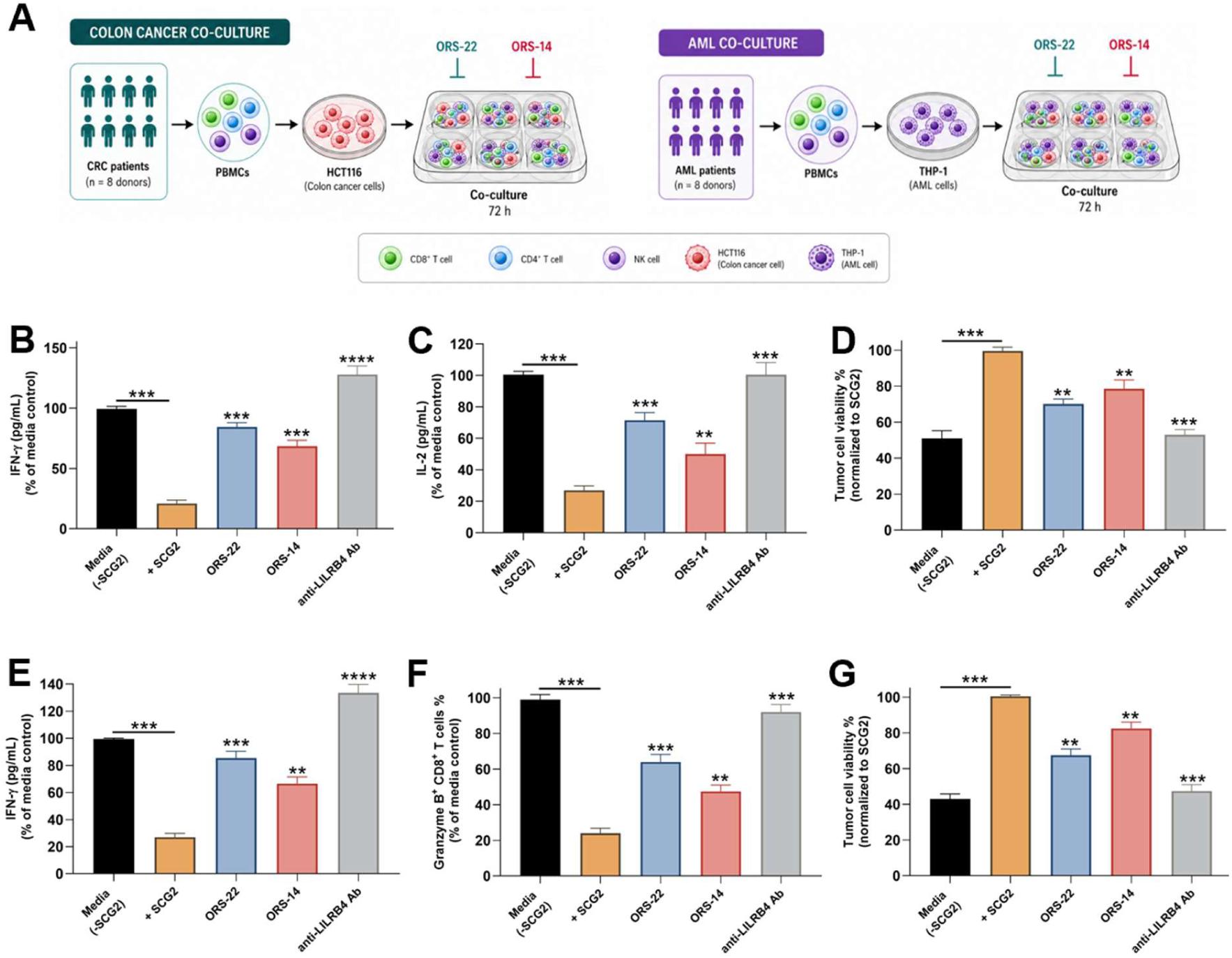
ORS-22 and ORS-14 restore anti-tumor immune activity across colorectal cancer and AML co-culture systems. **(A)** Schematic representation of ex vivo co-culture workflows used to evaluate the immunomodulatory activity of **ORS-22** and **ORS-14** in solid and hematologic tumor contexts. **(B,C)** In colorectal cancer co-cultures, SCG2 stimulation suppressed anti-tumor immune activity, resulting in reduced IFN-γ and IL-2 secretion. Treatment with **ORS-22** and **ORS-14** (1 μM) restored cytokine production, with **ORS-22** exhibiting greater activity. **(D) ORS-22** and **ORS-14** (1 μM) reduced HCT116 tumor cell viability in SCG2-treated CRC co-cultures, consistent with restoration of immune-mediated tumor killing. **(E)** In AML co-cultures, both compounds restored IFN-γ secretion suppressed by SCG2-mediated LILRB4 signaling. **(F) ORS-22** and **ORS-14** (1 μM) increased the frequency of Granzyme B^+^ CD8^+^ T cells, indicating recovery of cytotoxic T-cell activity. **(G)** Both compounds (1 μM) significantly reduced THP-1 AML cell viability in SCG2-treated co-cultures. In multiple functional assays, the activity of **ORS-22** approached that observed with a blocking anti-LILRB4 antibody (100 nM) used as a target-specific positive control. Data represent mean ± SD (n = 8 donors). Statistical significance was determined using one-way ANOVA followed by Tukey’s multiple comparisons test. \**p* < 0.05, \*\**p* < 0.01, \*\*\**p* < 0.001, \*\*\*\**p* < 0.0001.

The immunomodulatory activity of **ORS-22** and **ORS-14** further extended to hematologic tumor settings. In AML co-cultures containing patient-derived PBMCs and THP-1 AML cells, SCG2 stimulation similarly promoted immune suppression, resulting in reduced IFN-γ production, impaired cytotoxic T-cell activation, and increased AML cell viability. Both compounds significantly restored anti-tumor immune activity, as evidenced by enhanced IFN-γ secretion, increased frequencies of Granzyme B^+^ CD8^+^ T cells, and reduced AML cell viability (Figure 5E-G). These findings demonstrate that pharmacological blockade of LILRB4 using structurally distinct small molecules can reverse suppressive myeloid signaling across both solid and hematologic tumor contexts.

Importantly, the ability of **ORS-22** and **ORS-14** to phenocopy the effects of anti-LILRB4 antibody blockade across multiple orthogonal functional readouts strongly supports an on-target mechanism of action. The concordance between direct binding assays, CETSA target engagement studies, and downstream immune functional recovery provides a consistent mechanistic framework linking biochemical target engagement with restoration of anti-tumor immune responses. Collectively, these findings establish **ORS-22** and **ORS-14** as functional small-molecule modulators capable of reversing SCG2-driven immunosuppression and restoring productive immune activity in clinically relevant human tumor co-culture systems.

In summary, we have developed OracleScreen-LILRB4, a biophysical screening-trained ensemble machine learning framework that addresses the longstanding challenge of small molecule discovery against non-enzymatic immune checkpoint targets. By leveraging quantitative ΔF_norm_ binding signals from Dianthus-based TRIC screening of 19,800 compounds as a continuous regression target, OracleScreen-LILRB4 achieved strong predictive performance under scaffold-aware cross-validation and demonstrated exceptional prospective enrichment, identifying LILRB4 binders at a 15.0-fold higher rate than conventional screening. The identification of **ORS-22** and **ORS-14** as nanomolar-affinity LILRB4 binders with demonstrated cellular target engagement and functional immunomodulatory activity in patient-derived AML and colorectal cancer co-culture systems establishes a direct translational link between computational prioritization and functional small molecule discovery. The OracleScreen framework is generalizable and can be readily extended to other immune checkpoint targets for which biophysical screening data are available, offering a scalable and experimentally grounded approach to expanding the druggable space of the immune checkpoint landscape.

## Data Availability

The code is available on GitHub (DOI: https://github.com/gabr2003/OracleScreen-LILRB4.git).

## Author Contributions

The manuscript was written through contributions of all authors. All authors have given approval to the final version of the manuscript.

