## Supporting Information for "OracleScreen-LILRB4: Machine Learning-Guided Discovery of Myeloid Immune Checkpoint Binders Validated in Patient-Derived Cells"

**Table S1. SMILES strings and equilibrium dissociation constants (K_D_) of confirmed LILRB4 binders identified by OracleScreen-LILRB4 virtual screening.** All 16 compounds demonstrating concentration-dependent binding profiles in Dianthus TRIC dose-response screening are listed with their SMILES strings K_D_ values from nonlinear regression analysis of concentration-dependent ΔF_norm_ responses. Compounds are ranked in ascending order of K_D_ (highest to lowest affinity).

| **Compound number** | **SMILES** | **LILRB4 K_D_ from MST** |
| --- | --- | --- |
| **ORS-22** | CC1=NN=C2CN(CCN12)C(=O)CN(C)C3=NC=4C=CC=CC4S3 | 9.2 ± 0.6 nM |
| **ORS-14** | CC=1C=CC(=CC1C)OC=2C=C(C=CN2)CNC(=O)C3=CNN=N3 | 85.2 ± 11.3 nM |
| **ORS-37** | N#CC=1C=CC2=NC=C(CNC=3C=CC=C(C3)OC=4C=CC=CN4)N2C1 | 215 ± 27.4 nM |
| **ORS-2** | O=C(O)C=1C=CC(=CC1)OCC(=O)N2CCC(=N2)C=3C=CC=CC3 | 742 ± 81.3 nM |
| **ORS-7** | NC(=O)C1CCCN1CC=2C=CC=CC2NS(=O)(=O)C=3C=CC(=CC3)C=4C=CC=CC4 | 3.8 ± 0.62 μM |
| **ORS-13** | NC(=O)C1CC=2C=CC=CC2CN1CC3=NC(=NO3)C=4C=CC=CC4Cl | 7.7 ± 1.83 μM |
| **ORS-40** | CC1=CSC(=N1)N(C(=O)CNC(=O)C=2C=CC=CN2)C3CC3 | 10.2 ± 1.29 μM |
| **ORS-35** | O=S(=O)(C=1C=CC(=CC1)OCCN2C=NC=3C=CC=CC32)N4CCOCC4 | 15.9 ± 4.1 μM |
| **ORS-16** | C=1NN=C(C1CNCC=2C=CC=NC2)C3=CC=C4OCCOC4=C3 | 19.3 ± 3.02 μM |
| **ORS-3** | NC=1N=C(CN2C=CC=C(C2=O)C(F)(F)F)N=C(N1)NC=3C=CC=CC3 | 24.7 ± 6.41 μM |
| **ORS-8** | O=C(O)C1=CC=C(S1)C(=O)NC2CCCC3OCCC23 | 28.4 ± 5.82 μM |
| **ORS-17** | CC=1C=CN(N1)C=2C=CC(=CC2)C(=O)N3CCCC(C3)C(N)=O | 35.2 ± 4.24 μM |
| **ORS-44** | O=C(NC=1C=CC(=CC1)C2=CN3C=CSC3=N2)C=4C=CC(=CC4)S(=O)(=O)NCC5CCCO5 | 38.1 ± 6.31 μM |
| **ORS-51** | C=1C=CN2C(CC3=NN=C4C=CC=CN34)=NN=C2C1 | 41.3 ± 7.16 μM |
| **ORS-12** | NC(=O)C1CC=2C=CC=CC2CN1CCOC=3C=CC(=CC3)S(N)(=O)=O | 45.8 ± 2.46 μM |
| **ORS-55** | O=C(NC(CC=1C=CC=CC1)C(=O)N2CCNC(=O)C2)C=3C=CC(Cl)=CC3 | 52.5 ± 4.31 μM |

**Experimental methods**

**OracleScreen-LILRB4 Machine Learning Pipeline.** All computational analyses were performed in Google Colaboratory using Python 3.12. Molecular features were generated using RDKit (version 2026.3.2). For each compound, molecular representations were constructed by concatenating extended-connectivity Morgan fingerprints (radius = 2, 1024 bits) with ten physicochemical descriptors: molecular weight (MolWt), lipophilicity (MolLogP), hydrogen bond donor count (NumHDonors), hydrogen bond acceptor count (NumHAcceptors), topological polar surface area (TPSA), rotatable bond count (NumRotatableBonds), aromatic ring count (NumAromaticRings), fraction of sp3 carbons (FractionCSP3), heteroatom count (NumHeteroatoms), and ring count (RingCount). Compounds with invalid SMILES strings were excluded from analysis.

OracleScreen-LILRB4 was formulated as a regression task targeting continuous ΔF_norm_ binding signals rather than binary hit classifications, preserving the full quantitative information content of the Dianthus biophysical assay. Three ensemble algorithms were implemented: Random Forest (n_estimators = 300, min_samples_leaf = 2, class_weight = balanced), XGBoost (n_estimators = 300, learning_rate = 0.05, max_depth = 6), and Gradient Boosting (n_estimators = 200, learning_rate = 0.05, max_depth = 5). Final ensemble predictions were generated by averaging the predicted ΔF_norm_ scores across all three models.

Model performance was evaluated using scaffold-aware five-fold cross-validation. Murcko scaffolds were computed for all training compounds using RDKit, and compounds sharing identical scaffolds were assigned to the same cross-validation fold to prevent data leakage across structurally related series. Model performance was assessed using Spearman rank correlation coefficient between predicted and experimentally measured ΔF_norm_ values, as well as area under the receiver operating characteristic curve (ROC-AUC) using the continuous predicted score as a ranking signal for experimentally confirmed actives. Feature importance was assessed using SHAP (SHapley Additive exPlanations) analysis applied to the Random Forest component of the ensemble.

Virtual screening was performed by applying the trained OracleScreen-LILRB4 ensemble to a 45,760-member compound library. SMILES strings were processed identically to the training set, and ensemble-predicted ΔF_norm_ scores were generated for all compounds with valid SMILES representations. Compounds were ranked in descending order of ensemble score, and the top 200 predicted binders were selected for experimental validation by Dianthus TRIC screening. The code, datasets, virtual screening library, and top 200 compounds for experimental testing are available on Zenodo (DOI: https://doi.org/10.5281/zenodo.20350602).

**Dianthus TRIC Assay.** The Dianthus screening of chemical libraries for LILRB4 binding was performed using the temperature-related intensity change (TRIC) assay on a Dianthus NT.23 Pico instrument (NanoTemper Technologies). Recombinant human His-tagged LILRB4 protein (from SinoBiological, Catalog# 16742-H08H) was fluorescently labeled with RED-tris-NTA 2nd Generation dye (NanoTemper, Cat. #MO-L018) according to the manufacturer’s protocol.

Labeled LILRB4 (final concentration: 30 nM) was incubated with test compounds at a fixed concentration (10 µM) in assay buffer (PBS pH 7.4 supplemented with 0.05% Tween-20 and 1% (v/v) DMSO). Fluorescence was measured in Dianthus 384-well plates (NanoTemper Technologies, Catalog# DI-P001A) with LED power set to 40%. Fluorescence was recorded at 670 nm and 650 nm, and normalized fluorescence (F_norm_) was calculated as the ratio of F670/F650. Binding responses were quantified as Fnorm. Compounds with ΔF_norm_ ≥ 3 units and reproducible responses across replicates (n = 5) were classified as hits.

**Quantitative Binding Affinity Determination Using Monolith.** Binding affinities of selected hits were quantified by microscale thermophoresis using a Monolith X instrument (NanoTemper Technologies). His-tagged LILRB4 protein was labeled with RED-tris-NTA 2nd Generation dye using the Monolith His-Tag Labeling Kit (Cat. #MO-L018) following the manufacturer’s guidelines. Compound titrations were prepared as serial dilutions (PBS buffer, pH 7.4, 0.05% Tween-20, 1% (v/v) DMSO). Following a 15 min incubation at room temperature in the dark, samples were loaded into Monolith capillaries (Cat. #MO-K022) and analyzed at 25 °C using 40% LED power and medium MST power settings. Normalized fluorescence (F_norm_) values were determined as the ratio of fluorescence intensity after and before IR laser heating. Each compound was evaluated in five technical replicates. Dissociation constants (K_D_) were calculated from three independent experiments using MO.Affinity Analysis software and GraphPad Prism 10, applying standard dose-response fitting models. Data represent mean ± SD (n=5).

**CETSA.** CHO cells stably expressing human ILT3/LILRB4 (from Creative Biogene, Cat# CSC-RO0667) were maintained in Ham’s F-12 medium supplemented with 10% fetal bovine serum and 1% penicillin-streptomycin under standard humidified culture conditions (37 °C, 5% CO2). Cells were seeded in 384-well plates at 12,000 cells/well and incubated overnight at 37 °C in 5% CO₂. The tested compound was added at the indicated concentrations and incubated for 60 min. Cells were subsequently subjected to thermal challenge at 51 °C for 3 min (using a calibrated Bio-Rad C1000 Touch thermal cycler), followed by cooling to room temperature. Following thermal challenge, cells were lysed directly in assay plates by addition of lysis buffer supplemented with protease inhibitor cocktail and incubated for 30 min at 4 °C with gentle agitation. Lysates were subsequently clarified by centrifugation at 4000 × g for 10 min to remove aggregated and insoluble protein species generated during thermal denaturation.

The soluble fraction of LILRB4 remaining after thermal challenge was quantified using an AlphaLISA-based immunodetection format. Briefly, clarified lysates were incubated with LILRB4 -specific biotinylated detection antibody and AlphaLISA acceptor beads according to the manufacturer’s instructions, followed by addition of streptavidin-coated donor beads under reduced-light conditions. After incubation at room temperature, AlphaLISA signal was measured using Tecan Spark plate reader. CETSA stabilization signals were normalized to vehicle-treated controls, and dose-response curves were generated using nonlinear regression analysis. Data represent mean ± SD, n = 5.

**Human Co-Culture Assays.** For colorectal cancer co-culture assays, HCT116 cells (from ATCC) were seeded in flat-bottom 96-well plates at 1 × 10^4^ cells/well and allowed to adhere overnight in complete RPMI-1640 medium supplemented with 10% fetal bovine serum and 1% penicillin–streptomycin. PBMCs isolated from colorectal cancer patients (from STEMCELL Technologies) by Ficoll density-gradient centrifugation were added at an effector-to-target ratio of 10:1 in the presence or absence of recombinant human SCG2 (100 ng/mL) and 1 μM of each tested compound. A blocking anti-LILRB4 antibody (h128-3, catalog number HV126013, at 100 nM) was included as a target-specific positive control. Co-cultures were incubated for 72 h at 37 °C in 5% CO_2_. Supernatants were collected for IFN-γ and IL-2 quantification by ELISA (abcam catalog# ab46025 and #ab270883, respectively), and tumor-cell viability was assessed using CellTiter-Glo (from Promega) according to the manufacturer’s protocol.

For AML co-culture assays, THP-1 cells (from ATCC) were co-cultured with PBMCs from AML patients (from STEMCELL Technologies) or enriched CD3^+^ T cells under analogous conditions. Supernatants were collected for IFN-γ and IL-2 ELISA analysis (abcam catalog# ab46025 and #ab100706, respectively). Cytotoxic T-cell activation was assessed by measuring Granzyme B^+^ CD8^+^ T cells using flow cytometry. Tumor-cell viability was quantified using CellTiter-Glo (from Promega). Data were normalized to media or SCG2-treated controls as indicated in the figure legends. Data represent mean ± SD, n = 5.
